# Transcriptome and biochemical analysis pinpoint multi-layered molecular processes associated with iron deficiency tolerance in hexaploid wheat

**DOI:** 10.1101/2022.12.04.519030

**Authors:** Varsha Meena, Gazaldeep Kaur, Riya Joon, Anuj Shukla, Promila Choudhary, Palvinder Singh, Joy K Roy, Bhupinder Singh, Ajay K Pandey

## Abstract

Iron (Fe) is an essential nutrient for plants that is indispensable for many physiological activities. Although few genotypes were identified with contrasting tolerance to Fe deficiency, the molecular insight into the distinct biochemical and transcriptional responses determining the trait is poorly known. This study aimed to identify the molecular and biochemical basis for the contrasting Fe deficiency tolerance in wheat genotype showing tolerance to Fe deficiency (cv. Kanchan-KAN) compared to susceptible (cv. PBW343-PBW) cultivar. Under Fe deficiency, the KAN show delayed chlorosis, high SPAD values and low malondialdehyde activity compared to PBW. The shoot transcriptomics studies show that a large set of genes for photosynthetic pathways were highly induced in PBW, suggesting its sensitivity to Fe deficiency. Although, under Fe deficiency, both the cultivars show distinct molecular re-arrangements, including high expression of genes involved in Fe uptake (including membrane transporters) and mobilization, the gene expression level was higher in KAN. Furthermore, the KAN cultivar also shows high ubiquitination activity in the shoot tissue suggesting a high turnover of proteins in the tolerant cultivar. These observations were also co-related with the high root phytosiderophores biosynthesis and its release that contributes to the enhanced Fe translocation index in KAN. Overall, our work provides the key link to understanding the mechanistic insight for the Fe deficiency tolerance in hexaploid wheat. This will enable wheat breeders to select genotypes for better Fe use efficiency for agriculture.

## Introduction

Iron (Fe) is an important nutrient for plants, as it is required in many vital functions ranging from photosynthesis to respiration and other essential cellular function. Fe plays a crucial role in plant functions that require electron transfer reactions, including photosynthesis and nitrogen assimilation. Although Fe is abundant in soil, its bioavailability is restricted at high pH, particularly in aerobic and calcareous soils. Alkalinity in soils, mainly due to bicarbonate (HCO3^-^) ions, leads to the reduced solubility of Fe oxides in soil [1,2]. Earth’s 30% soil is alkaline and, thus, has low bioavailable Fe that could cause severe Fe deficiency in crop plants. This deficiency leads to decreased crop yield and nutritional quality of the grains. To cope with this, plants have developed strategies to combat micronutrient deficiency by recruiting specific molecular components [3]. These components/strategies differ depending on the type of plant species, mono or dicots. Dicots generally use a reduction-based strategy (Strategy-I), while monocots use a chelation-based strategy (Strategy II). Strategy I mainly relies on two ways: the P-type ATPase helps enhance the proton extrusion to decrease the pH [4], and plasma membrane-bound reductase at the root surface induces the reduction of Fe^3+^ to Fe^2+^ [5]. Graminaceous species, including wheat, depict strategy II to improve Fe acquisition. These plants release certain chelating compounds known as phytosiderophores (PS) or mugineic acids (MAs) via the PS efflux transporter [6]. These PS chelate metal ions and help in the uptake of Fe and other micronutrients root via high-affinity transporter YSL [7,8].

To reduce the agricultural and economic impact of Fe deficiency stress, it has become necessary to develop a Fe-efficient plant capable of optimum use of Fe for its metabolism. Some plants can have the same high yield by just a smaller amount of Fe uptake than less efficient plants [9,10]. Selecting the Fe efficient plants could help in limiting the use of Fe fertilizers, thus curbing the economic losses that arise due to Fe deficiency. Fe deficiency tolerance among the species of graminaceous plants was found to be linked with the amount, and types of MAs secreted [11,12]. Genotypes with efficient micronutrient acquisition are considered better adaptable to deficient soils. Sufficient genetic variability exists among the different genotypes with differential nutrient acquisition and utilization efficiency in low micronutrient environments [13,14]. Additionally, Fe limitation in the rhizosphere could alter the molecular and biochemical process specific to shoots. Typically, a delayed chlorotic response in shoots is considered an important trait that helps minimize the micronutrient deficiency impact. Limited attempts have been made to identify the Indian wheat cultivars with tolerance to Fe deficiency condition, but no molecular insight was provided for this tolerance trait [15]. An insight into the available literature reveals that the information about varying shoot Fe demand is transmitted through signals between shoot and root, and variation in Fe absorption, transport and storage is observed [16,17].

Wheat is an important crop (*Triticum aestivum* L.) and the most widely grown cereal crop accounting for a total of 20% of calorific intake by humans. Previously, molecular signals induced during Fe deficiency were studied in roots [18,19]. These studies demonstrated the importance of the root molecular components involved in Fe uptake and mobilization as a response to Fe deficiency. However, how the genes in shoots are affected by Fe deficiency remains largely unaddressed. Recently, it was shown that Fe status in the shoot and the hormone including auxin largely induce the PS release in the roots [11]. Therefore, characterizing the shoot molecular response could activate and regulate the Fe deficiency response in roots. Alternatively, utilizing the contrasting wheat cultivars for their Fe deficiency response could provide much-needed insight and delineate pathway that contributes to Fe deficiency tolerance [15].

In the current work, morpho-physiological and biochemical characteristics, including high PS release, were observed to confirm the contrasting tolerance to Fe deficiency among wheat cultivars. At the molecular level, shoot RNAseq analysis revealed the distinct molecular changes contributing to the delayed chlorosis and multiple pathway genes that contribute to the Fe deficiency tolerance. This study provides insight into our understanding of the molecular cross-talk required to develop the Fe deficiency tolerance trait that could help minimise the loss in crop productivity in nutrient-deficient soils.

## Materials and methods

### Plant materials and experimental setup

Wheat (*Triticum aestivum)* cv. Kanchan (KAN) and PBW343 (PBW) were used in this study (obtained from Indian Agricultural Research Institute, Delhi). Seeds were thoroughly rinsed with autoclaved MQ followed by surface sterilization with 1.2% sodium hypochlorite and 10% ethanol for 3 min. Seeds were rinsed 2-3 times with autoclaved MQ and kept at 4°C overnight for stratification and then allowed them to germinate on petri dishes with 2-3 layer of wet Whatman paper for 5-7 days. Once leafy stage was reached, the endosperm was removed, and uniform seedlings were subjected to hydroponic system consist of PhytaBox™ containing Hoagland nutrient solution. For control, following composition of Hoagland media was used 6 mM KNO3, 1 mM MgSO_4_, 2 mM Ca(NO_3_), 2 mM NH_4_H_2_2PO_4_, 80 μM Fe-EDTA, 25 μM H_3_BO_3_, 2 μM MnSO_4_, 0.5 μM CuSO_4_, 2 μM ZnSO_4_, 50 μM KCl and 0.5 μM Na2MoO_4_. For Fe starvation condition 1 μM Fe-EDTA was used. Seedlings were grown in aerated condition, and media was changed at every third day to avoid any nutrient depletion. Each experiment had three biological replicates. Sample collection was done at three-time points 8, 11 and 14 (DAT) day after treatment. Based on distinct phenotypes, shoot samples collected at 8 days were used for transcriptome analysis with three replicates of each treatment. All the seedlings were grown in a growth chamber under controlled environmental conditions at 22–24 °C temperature, 65%–70% humidity, 16 h of photoperiod along with photosynthetically active radiation of 300 μmol m^−2^s^−1^ of light.

### RNA extraction and qRT-PCR analysis

Shoot samples collected at 8 days after Fe starvation were finely ground in liquid nitrogen. Total RNA was isolated from 100mg of grounded tissue using Spectrum plant total RNA kit (Sigma), following the DNAse treatment using Turbo DNAfree kit (Invitrogen, USA). 2 μg of total RNA from each sample was used for cDNA synthesis using SuperScript III First-Strand Synthesis System (Invitrogen, USA as per manufacturer’s instructions). Expression analysis of wheat transcripts in different genotypes was quantified by PCR system (Biorad, USA) and 1:10 dilution of cDNA was used to quantify genes using SYBR Green I (QuantiFast® SYBR® Green PCR Kit, Qiagen, USA). *ADP*-*Ribosylation Factor 1* (AB050957.1) was used as the internal reference to normalize the gene expression. Fold changes of gene expression were calculated using delta-CT method (2^−^_ΔΔ_CT) [20].

### Malondialdehyde estimation and Ferroxidase activity

Malondialdehyde (MDA) level was carried out as described previously [21]. Briefly, 100 mg of leaves tissue was homogenized in 0.1% TCA followed by centrifugation at 4 °C with 13000 rpm for 10 min. 400 μl of supernatant was added to 1ml of 0.5 % TBA in 20 % TCA and incubated at ice for 5 min followed by centrifugation at 13500 rpm for 5min. The absorption of the supernatant was taken at OD 532 and 600 nm. MDA was estimated by using the extinction coefficient of 155 mM^-1^ cm^-1^.

The root ferroxidase was measured as the oxidation of Fe^2+^ form, which was assayed by monitoring the rate of disappearance of ferrous ammonium sulphate using the ferrous chelator ferrozine. An equal amount of total protein (15 μl of total protein (2 μg/ml) from the roots subjected to Fe deficiency condition was used to perform a reaction containing 1050 μl buffer (450 mM Na-acetate of pH 5.8), 100 μl CuSO_4_) by initiating the reaction with addition of 225 μl of the substrate (Fe(II)SO_4_, 357μM) containing 100 μM CuSO_4_ [22]. The aliquots (200 μl) were then removed at appropriate intervals and transferred to 96 well microtiter plates for reaction-quenching with 14 μl of 18 mM ferrozine. The rate of Fe^2+^ oxidation for the Fe^2+^-ferrozine complex was calculated from the decreased absorbance at 560nm, and the graph was plotted [23]. These experiments were repeated three to four times, each containing eight to ten wheat seedlings.

### Elemental analysis using ICPMS

Metal analysis was performed as described earlier [24]. Briefly, root and shoot tissue was finely ground and incubated at 60°C until the constant weight reached. 100 mg of dry tissue were digested using HNO_3_ (SuraPure™, Merck) in the microwave. Digested samples were subjected to mass analysis using the ICPMS system. Three technical replicates were used for metal analysis for these three biological replicates.

### Estimation of total PS

PS release was measured by estimating its content in the root washing as previously described [25]. After 2h of onset of light 10 seedlings per treatment were washed with deionised water for 1 min then were submerged in 20 ml of double deionised aerated water for 4h under dark conditions. A fresh solution of Fe(OH)_3_ was prepared by adding 1N NaOH in 4Mm FeCl_3._ In the above root washing, 2ml Fe(OH) and 0.5ml of 0.5 M Na-acetate buffer (pH 5.6) were added and kept on a shaker for 30 min. This solution was then filtered into the flask containing 0.2ml 6N HCL using whatman#1 filter paper, followed by the addition of 8% hydroxylamine-hydrochloride and kept at 60 °C for 20 min. To it, 0.2 ml of 0.25% of ferrozine and 1ml of 2M Na-acetate buffer (pH 4.7) were added. Total PS was measured by estimating reduced Fe in solution by taking OD at 562nm.

### In Vivo Ubiquitination Analyses

In vivo ubiquitination analysis was performed as [26] described with the following modifications. A total of 10 seedlings for each sample were ground in liquid nitrogen and then homogenized with protein extraction buffer containing 50 mM Tris (pH 8.0), 150 mM NaCl, 1% Triton X-100, 1 mM EDTA, 10% glycerol, and 1× phenylmethylsulfonyl fluoride (PMSF). Total protein concentration was measured, and a 100 μg protein sample was separated on a 10% (w/v) PAGE and visualized by staining with Coomassie Brilliant Blue or immunoblotted with anti-Ub (Invitrogen; pa1-10023).

### RNAseq and Differential Expression Analysis

RNA was isolated manually in triplicates using TRIZOL method. Total RNA quality and quantity were checked using nanodrop and gel electrophoresis in 1% denaturing RNA agarose gel. RNA samples (12 RNA samples representing three experimental replicates for each genotype) with RNA Integrity Number (RIN) greater than eight were used for library preparation and sequencing. Transcriptome sequencing was performed using Illumina HiSeq1000 sequencing technology and deposited at NCBI (Submission: SUB12344025 with BioProject: PRJNA907484).

Transcriptome analysis was performed, as mentioned earlier [18]. In summary, Trimmomatic 0.35 was used for clipping and quality-trimming the reads. We removed the reads with a Phred scaled quality score of less than Q20. After removing adapter sequences and filtering the low-quality reads, they were mapped to the reference genome using TopHat v2.1.1 with default parameters. The Cufflinks v2.2.1 program assembled the transcriptome data from RNA-seq data (Submission: SUB12344025 with BioProject: PRJNA907484and was also used to quantify their expression. Mapped reads were subjected to Cufflinks, followed by Cuffmerge and Cuffdiff. Log2 fold change (FC) values >1 were considered up-regulated, whereas those with an FC <1 were considered down-regulated. These genes were further categorized based on statistical significance (*P*<0.05) and the false discovery rate (FDR 0.05) after Benjamin–Hochberg corrections for multiple testing for their significant expression. For each genotype, DEGs were identified with respect to their control treatments. The data generated from this study have been deposited in the NCBI Sequence Read Archive (SRA) database and are accessible with the submission ID-SUBxxxx and BioProjectID-xxxx.

## Results

### Fe deficiency tolerance response in KAN and PBW cultivars

Previously, wheat cultivars screening showed varying Fe deficiency tolerance [27]. We selected two wheat cv. KAN (tolerant) and PBW (susceptible) show contrasting phenotypes to unravel the molecular basis of Fe deficiency tolerance. Wheat seedlings subjected to Fe deficiency condition were monitored, and on the 8^th^ day of its growth, intense chlorotic symptoms on the young leaves were observed in PBW compared to KAN (Figure 1A). SPAD index shows a drastic reduction in PBW leaves than in KAN cultivars compared to their respective controls suggesting their sensitivity towards Fe deficiency (Figure 1B). Interestingly, Fe deficiency affects the root growth in both genotypes, pointing to a general symptomatic response to this nutrient. Under control conditions, no significant differences in the root lengths were observed in the genotypes, but in Fe deficiency, the percentage reduction in the root length was higher in PBW (Figure 1C). Plants under stress could show lipid peroxidation that could help in regulating the Fe deficiency stress in roots. In our experiments, both genotypes show similar net increases in the MDA accumulation compared to the respective control (Figure 1D). These biochemical and physiological observations suggest that PBW is highly sensitive and KAN is tolerant to Fe deficiency.

**Figure 1:**
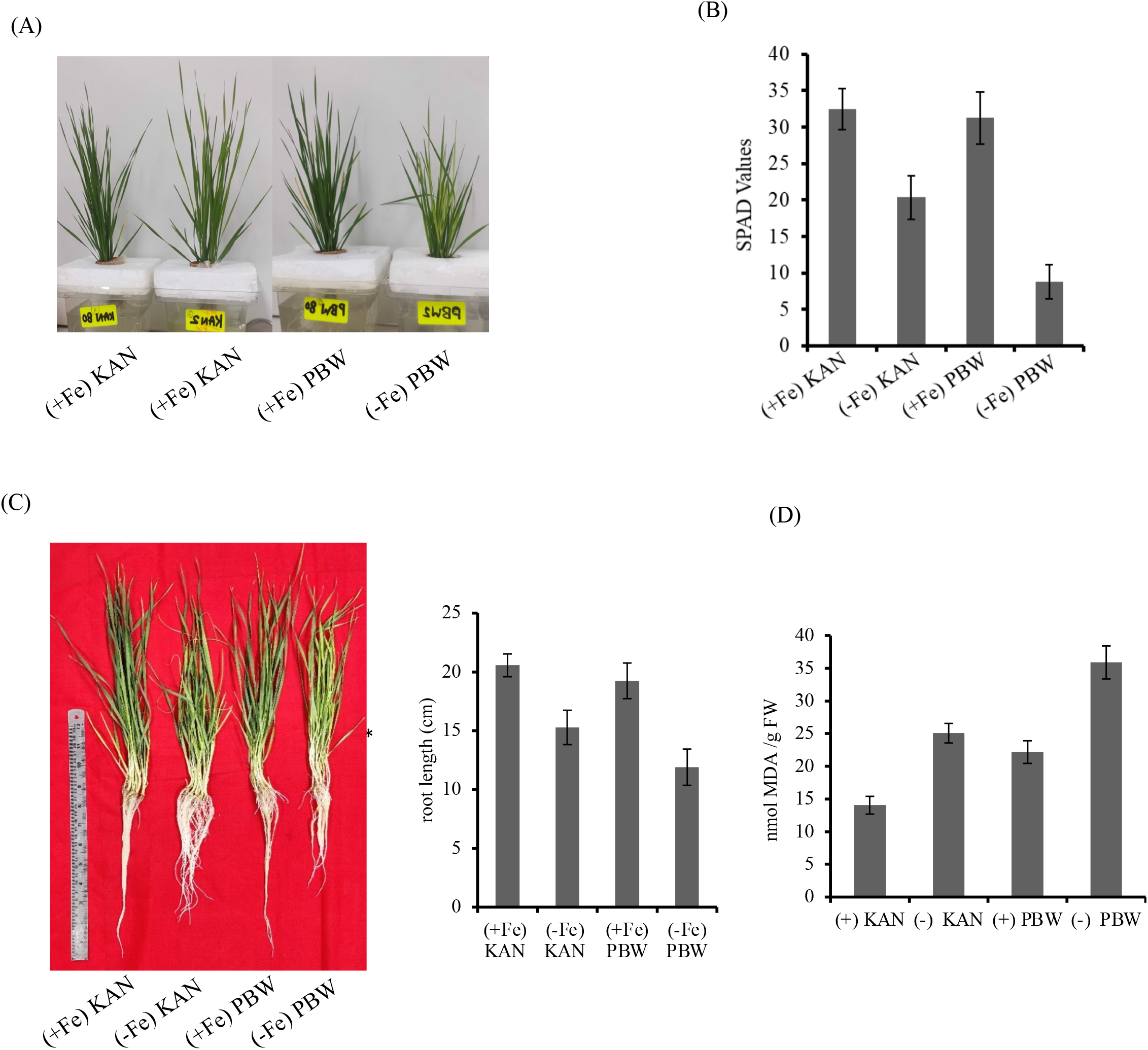
Effect of Fe deficiency on the plant growth of two hexaploid wheat cultivars KAN and PBW. (A) Phenotypes of KAN and PBW under Fe deficiency condition (2uM Fe-EDTA). (B) Leaf chlorophyll level (SPAD index) of the wheat cultivars under Fe deficiency and control condition. Presented values are mean±SD, n=8. (C) Root length (cm) for the two genotypes as observed. Presented values are mean±SD, n=8. (D) Estimation of the MDA content in the shoots of wheat cultivars. Presented values are mean±SD, n=8.

### Shoot transcriptional response of wheat to Fe deficiency stress

We performed transcriptome analysis for the wheat shoots under Fe deficiency conditions to account for the molecular changes in the contrasting wheat genotype. Our three biological replicates reveal the consistency in the replicates as projected in the PCA plot (Supplementary Figure S1). The generated reads were mapped to the Wheat Ensembl Plants Release-41 reference genome that resulted in providing the mapping % reads ranging from 85.57 to 96.01 %. Our analysis revealed 4908 and 5579 genes differentially expressed in KAN and PBW genotypes compared to their respective controls. Out of these, a total of 2588 and 2912 genes were upregulated in KAN and PBW, whilst 2320 and 2667 genes were downregulated in these genotypes (Figure 2A). When comparison was drawn among the genotype under the Fe deficiency, total 10,679 DEGs were identified out of these 5957 were upregulated and 4722 found to be downregulated. These observations suggest that a distinct genotypic response to Fe deficiency that was found to be characteristic for Fe-stress.

**Figure 2:**
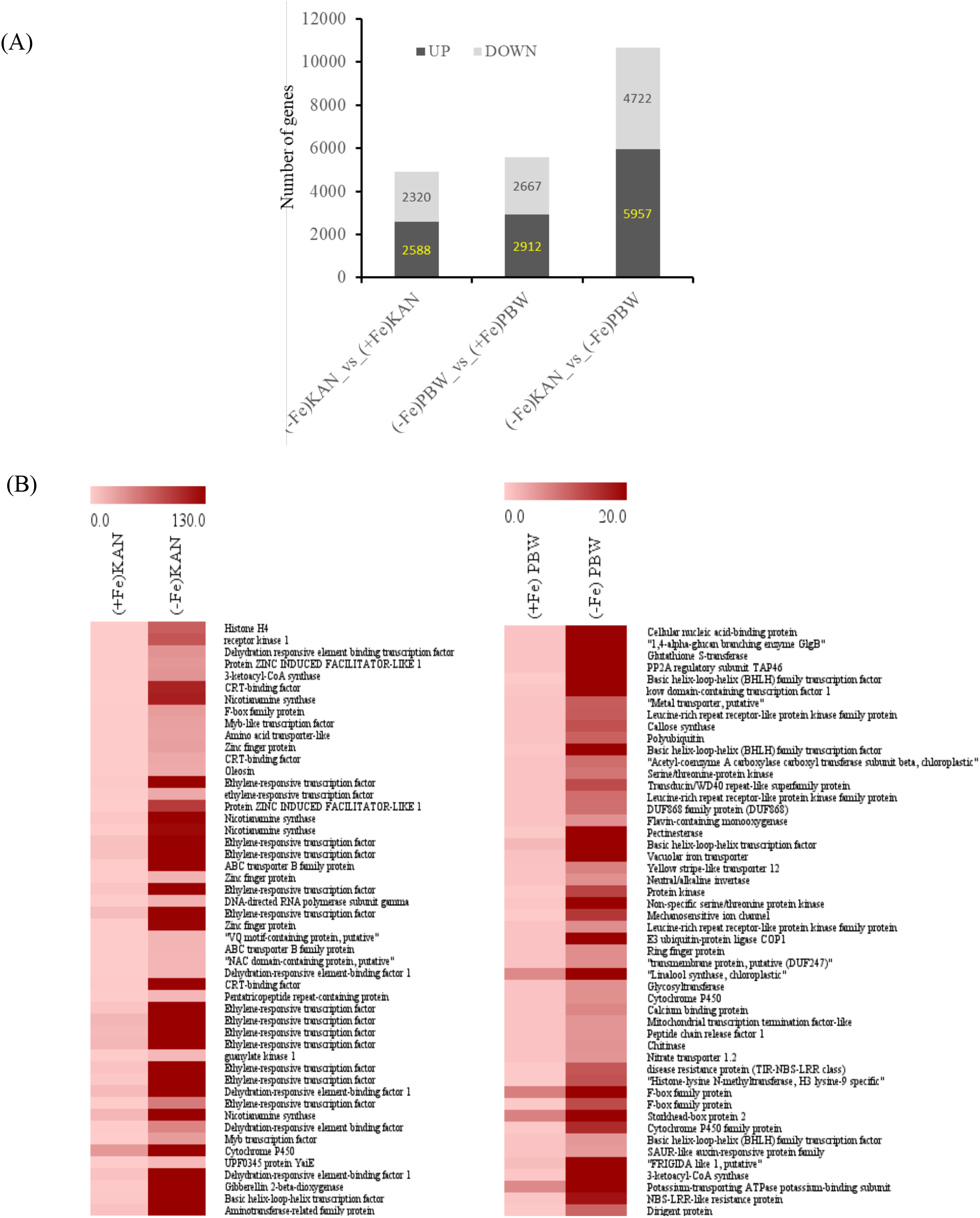
Differential expressed genes (DEGs) in the shoots of KAN and PBW under Fe deficiency. (A) Bar indicating the number of DEGs expressed in the shoot tissue of the wheat cultivars. (B) Heatmap analysis of top fifty most differentially expressed genes in KAN and PBW cultivars. Light brown to dark brown colour indicates the intensity of the LOG2FC changes in the gene expression with respect to their controls.

### Distinct molecular response of the wheat cultivars to Fe-deficiency

We performed the heat-map analysis of the top 50 genes that were differentially regulated under the Fe deficiency in both genotypes and their respective controls. Our heat maps indicated the expression of multiple gene family genes at very high fold levels. Within the top 50 DEGs for each cultivar, we see a different dataset pointing to the distinct gene families. For example, KAN show multiple genes involved in strategy-II mode of Fe uptake, mobilization and TFs (i.e. AP2/ERF) and dehydration-responsive element binding factor-1 (DREB) dehydration-related genes encoding genes (Figure 2B). In contrast, the top 50 genes in PBW include transcripts encoding for metabolic pathways such as cytochrome-P450, phenylpropanoid pathway (PPP) genes and photosynthesis-related genes.

To get insight into how the DEGs represent metabolic pathways and other changes, we compared the datasets for each genotype with their control and between the genotype. Our analysis reveals distinct metabolic pathway genes perturbed post-Fe deficiency in these genotypes. The data revealed that PBW show more differential regulation of the genes for pathways such as secondary metabolism, lipid biosynthesis related and light reactions compared to control (Figure 3A). In contrast, low genes were represented for the above-mentioned pathways in KAN grown under Fe deficiency compared to control (Figure 3A). Interestingly, when a comparison was drawn amongst the cultivar for the DEGs, a large number of common pathway genes were perturbed in KAN compared to PBW for the pathways such as secondary metabolism, amino acid biosynthesis, light reactions, ascorbate and glutathione biosynthesis, sugar metabolism and cell wall biosynthesis genes (Figure 3B). Furthermore, the bubble analysis also suggested the high enrichment of these specific pathway genes (Figure 3C). Based on these observations, we conclude that KAN seedlings could adapt better to the Fe-deficient stress than the PBW.

**Figure 3:**
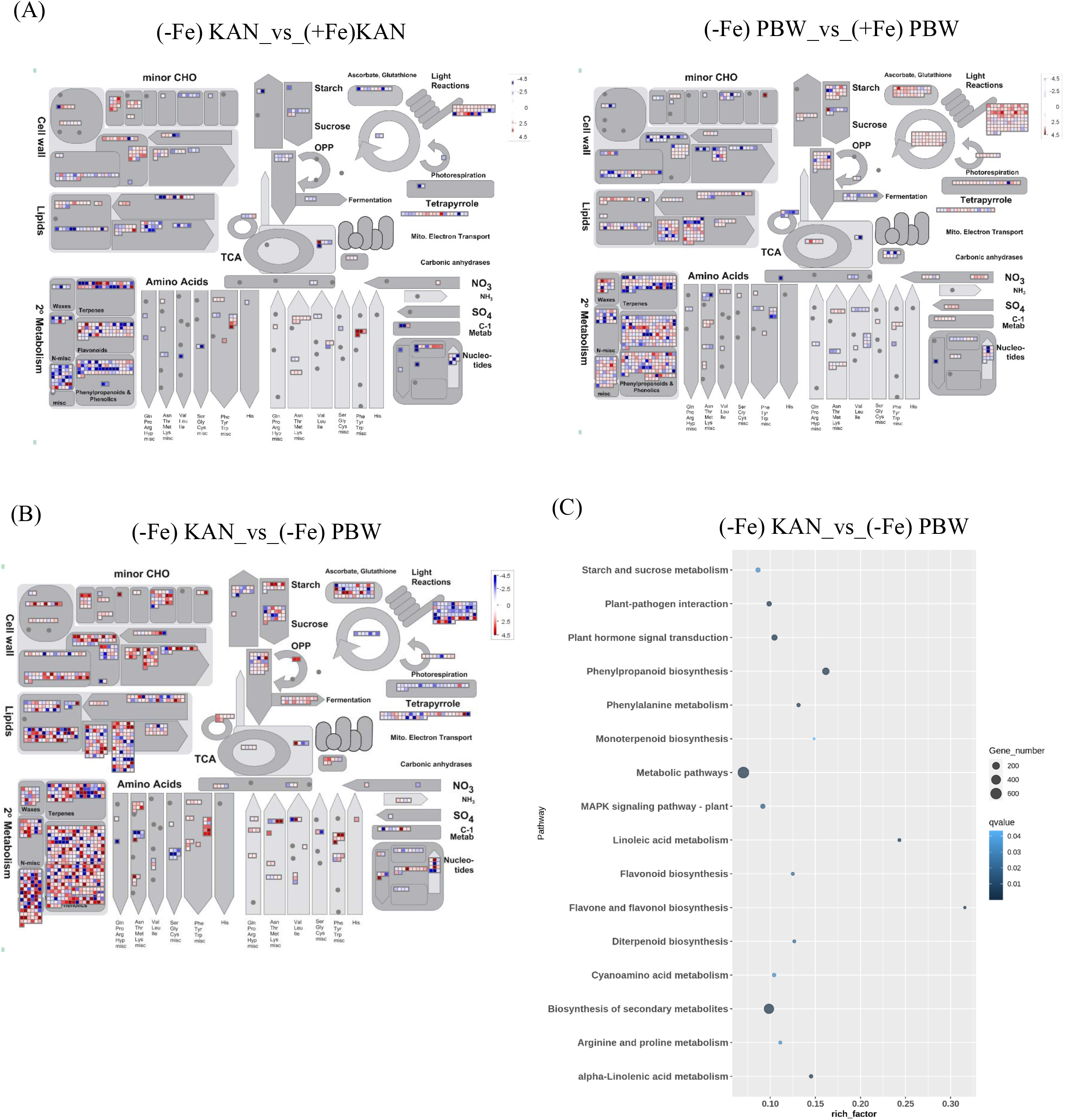
Map-man and gene enrichment analysis of the DEGs in the KAN and PBW cultivars under Fe deficiency. (A) Mapman analysis of the respective DEGs in the Kan and PBW cultivars under Fe deficiency conditions. (B) Mapman analysis of the respective DEGs under Fe deficiency in the Kan cultivar with respect to PBW. (C) KEGG pathway enrichment analysis for Fe deficiency response in KAN cultivar with respect to the PBW. Enrichment pathways (qvalue<0.5) for KAN with respect to PBW. X-axis indicates the rich-factor of the perturbed respective pathway genes with respect to the total number of genes involved. Bubble size indicates the number of genes altered in the respective terms.

### PBW show aggravated Fe deficiency-induced photosynthesis pathway genes

Photosynthetic activity is highly dependent on the presence of Fe. We checked how photosynthesis antenna proteins are affected in Fe deficiency conditions in these genotypes compared to the control. We focused on the known genes involved in the photosynthesis photosystems (Figure 4A). Our analysis shows 146 DEGs in PBW and only 35 genes in KAN were differentially expressed with respect to their control (Table S1). Subsequently, heatmaps for a sub-set of genes were prepared for the DEG with logFC>1.0. This analysis suggests Fe deficiency could impart changes in the photosynthesis-related genes more strongly in PBW. Specifically, in this genotype, genes involved in photosystem-I, photosystem-II, ferredoxin and ATP synthase were highly affected (Figure 4B and Table S1). Interestingly, both genotype shows limited changes in the genes involved in the state transition. This suggests that although molecular remodelling of the photosynthesis super-complexes shows similar changes, high differential changes related to other photosystem genes were observed in PBW compared to KAN. The molecular changes in the shoots could transduce signals in the roots. Therefore, it will be interesting to study the role of transcription factors that could affect the early chlorotic response and control the overall molecular changes in both genotypes.

**Figure 4:**
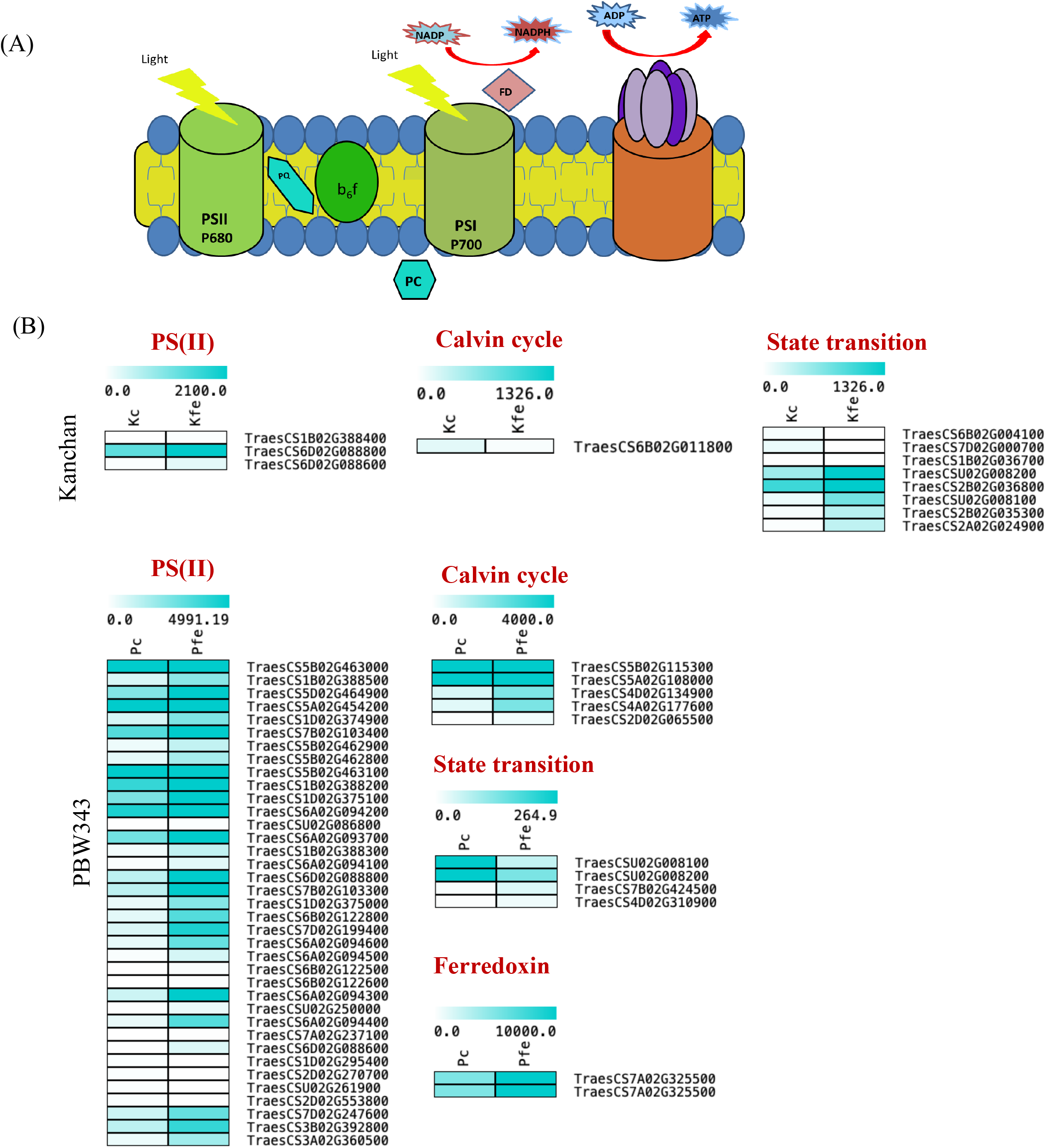
Effect of Fe deficiency on the photosynthesis pathway genes. (A) Schematic representation of the photosynthesis pathway in plants. (B) Heat map analysis of the pathway genes involved in the different stages of photosynthesis in KAN and PBW under Fe deficiency.

### High regulatory activity in KAN compared to PBW

Next, we checked if there is any unique involvement of transcription factors in the respective cultivars. Earlier, bHLH, AP2/ERF, MYB, WRKY and Dof TFs were expressed differently in Fe deficiency [28]. To our surprise, we observed a high number of TFs induced in the KAN cultivar compared to PBW. The dominant genes encoding for the TFs, including WRKY, bHLH, ERTF and MYB were abundant in KAN and PBW (Figure 5A). The 10.71% of genes encoding for TFs in KAN accounted for the total DEGs; whereas only 7.29 % of TFs represent DEGs in PBW (Table S2 and S3). This striking observation suggests that TFs plays a major role in imparting the Fe deficiency tolerance response in KAN. The high expression burst of TFs in tolerant cultivar signifies quick and regulated responses to manage the Fe deficiency condition.

**Figure 5:**
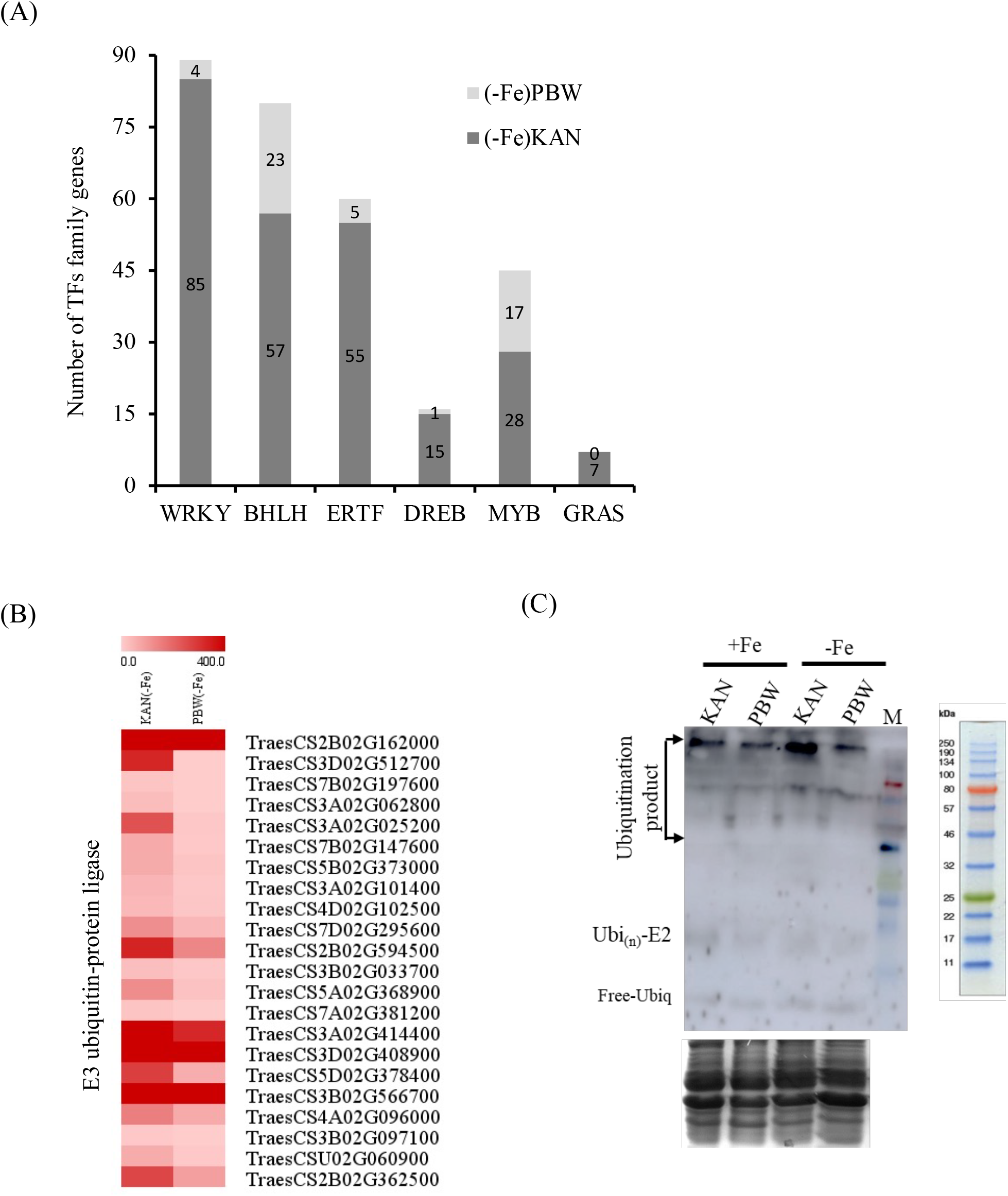
Expression analysis of genes encoding for the transcription factors (TFs) and E3 ubiquitin ligase. (A) Number of TFs those were expressed in KAN and PBW under Fe deficiency with respect their control treatment. (B) Heatmap analysis of genes encoding for the E3 ubiquitin ligase in KAN and PBW. (C) Western analysis for the Ubiquitination assay in KAN and PBW cultivars subjected to Fe deficient (-Fe) and sufficient condition (+Fe).

Earlier, we observed many photosynthesis-related genes being differentially expressed in PBW. Therefore, we analysed the promoter region of genes (LFC>1.5) involved in the photosynthesis process. Interestingly, our analysis revealed a high number of the cis-element region for AP2/ERF, bHLH, MADS, WRKY, bZIP binding proteins (Table 1). This suggests that in addition to regulating genes for Fe-uptake and translocation, multiple TFs could regulate photosynthetic-related activities.

**Table 1:**
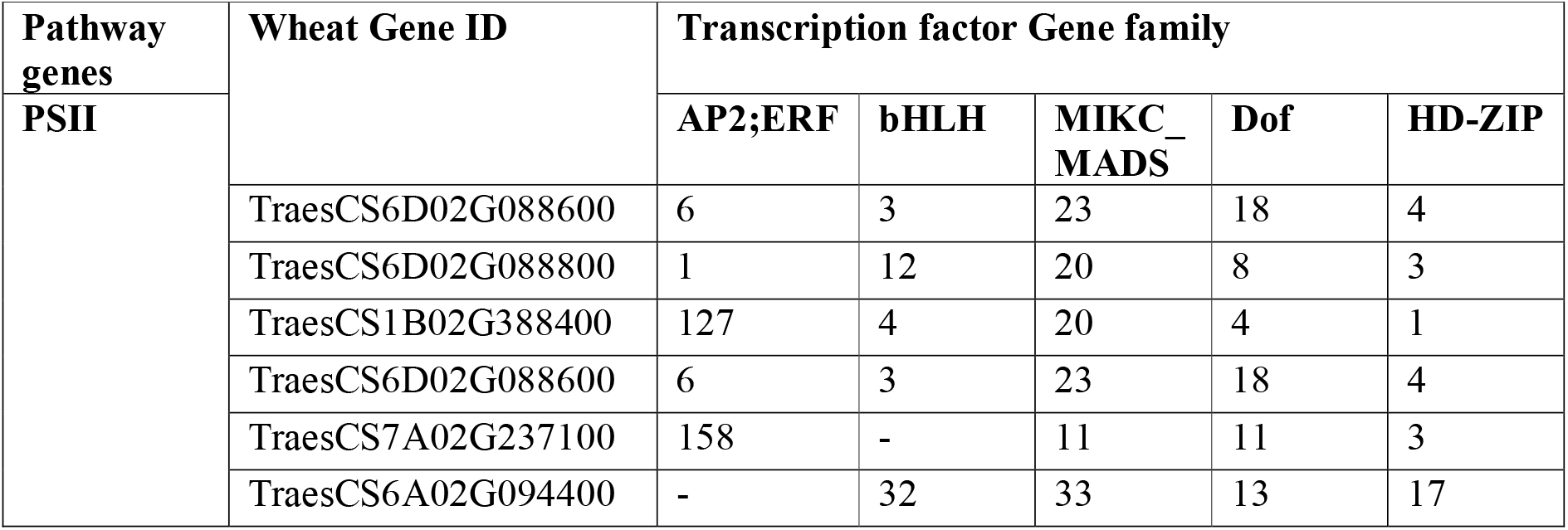
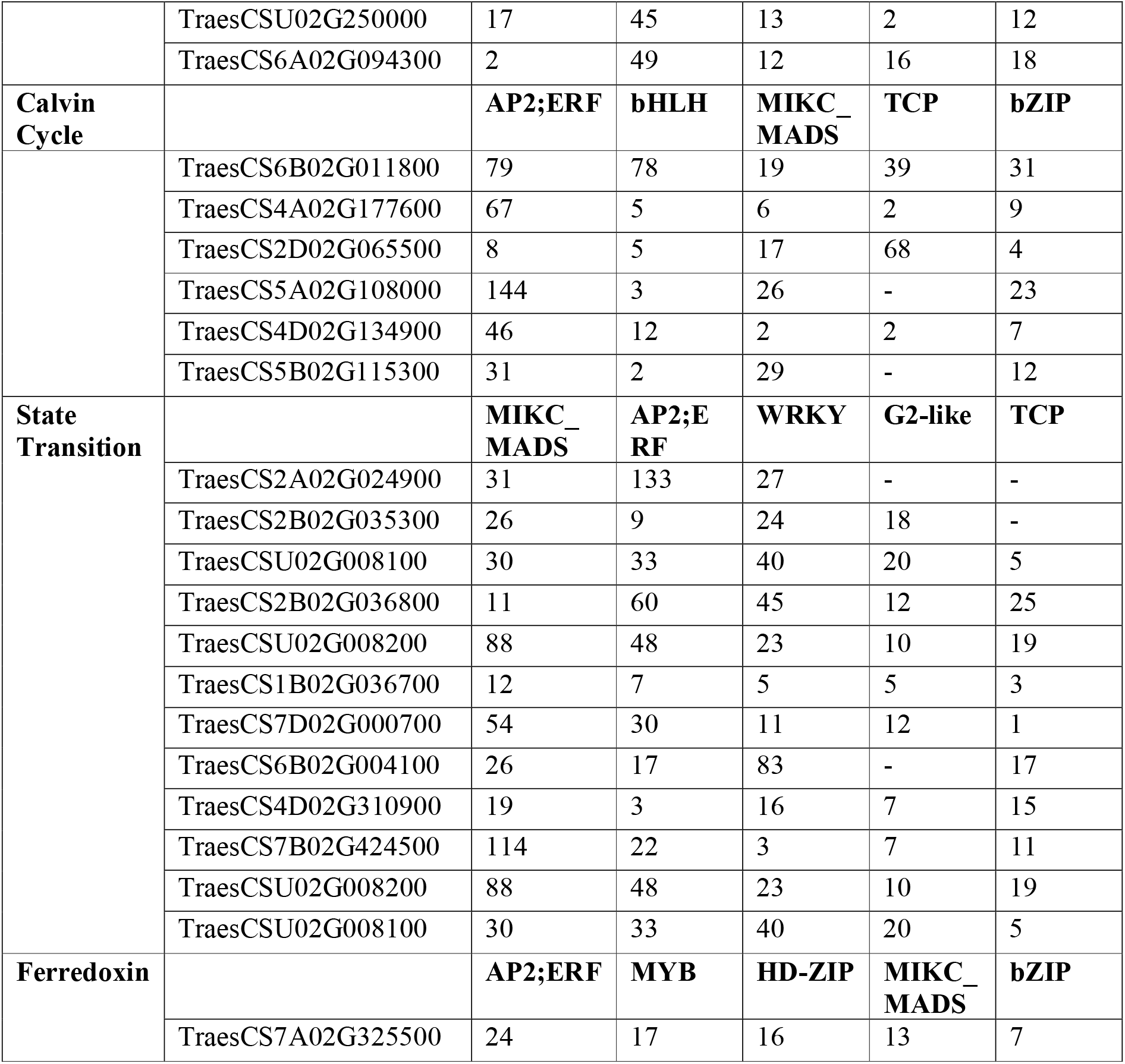
Putative motifs predicted in promoters of photosynthesis related genes differentially regulated (LFC>1.5) during Fe-deficiency across the wheat genotype. Transcription factor binding site counts were extracted using fimo tool and in-house perl scripts for each gene.

We were intrigued by the abundance of genes encoding the E3 ubiquitin-protein ligase among the DEGs. Previously, the role of Ubiquitination mediated by E3 Ubiquitin ligase has been demonstrated in controlling the turnover of metal transporters and TFs [29–31]. In the present study, multiple E3 ligases were differentially regulated (Supplementary Table S4). Interestingly, as many as 18 E3 ubiquitin ligases were upregulated in KAN genotype under Fe deficiency in contrast to PBW (Figure 5B). For this point of Fe deficiency, the upregulation of E3 ligases seems very specific in the cv. KAN. Next, we performed the Western analysis to check if the differential expression of the E3 Ubiquitin ligases could affect the ubiquitinated products in these contrasting cultivars. Our data suggest the formation of more ubiquitinated products in KAN that were captured using ubiquitin antibody as compared to the PBW (Figure 5C). Our analysis indicates that the robust response of TFs and ubiquitination could regulate the protein turnover at the post-transcriptional and translational levels to impart Fe deficiency tolerance.

### KAN show high PS release and surge in the genes for Fe mobilization to shoots

To account for the enhanced tolerance level due to the molecular changes of the DEGs we then measured the Fe translocation index, ferroxidase activity, and the PS release ability of these two genotypes post-Fe deficiency. We measured the content of the Fe in the shoots and roots to calculate the translocation index of Fe. Our result shows a high translocation of Fe in the KAN genotype compared to the PBW. At this point, the mean value of the translocation index for KAN was 0.35 (mean value) compared to the PBW with 0.26 (Figure 6A). The high translocation index of Fe was speculated due to the enhanced availability of Fe in the rhizosphere. It could be due to the expression of genes involved in Fe-homeostasis [32–34]. Earlier the PS release has been linked with the Fe deficiency tolerance in different crops species [15]. Interestingly, both cultivars show similar ferroxidase activity in the roots (Figure 6B). Next, we measured the total PS released by these genotypes under Fe deficiency. As expected, the Fe-sufficient plants show very low levels of PS compared to Fe-deficient plants. We observed that under the Fe deficiency, the roots of KAN release a high amount of PS (∼58 nmMol) when compared to the PBW (8 nmMol) (Figure 6C).

**Figure 6:**
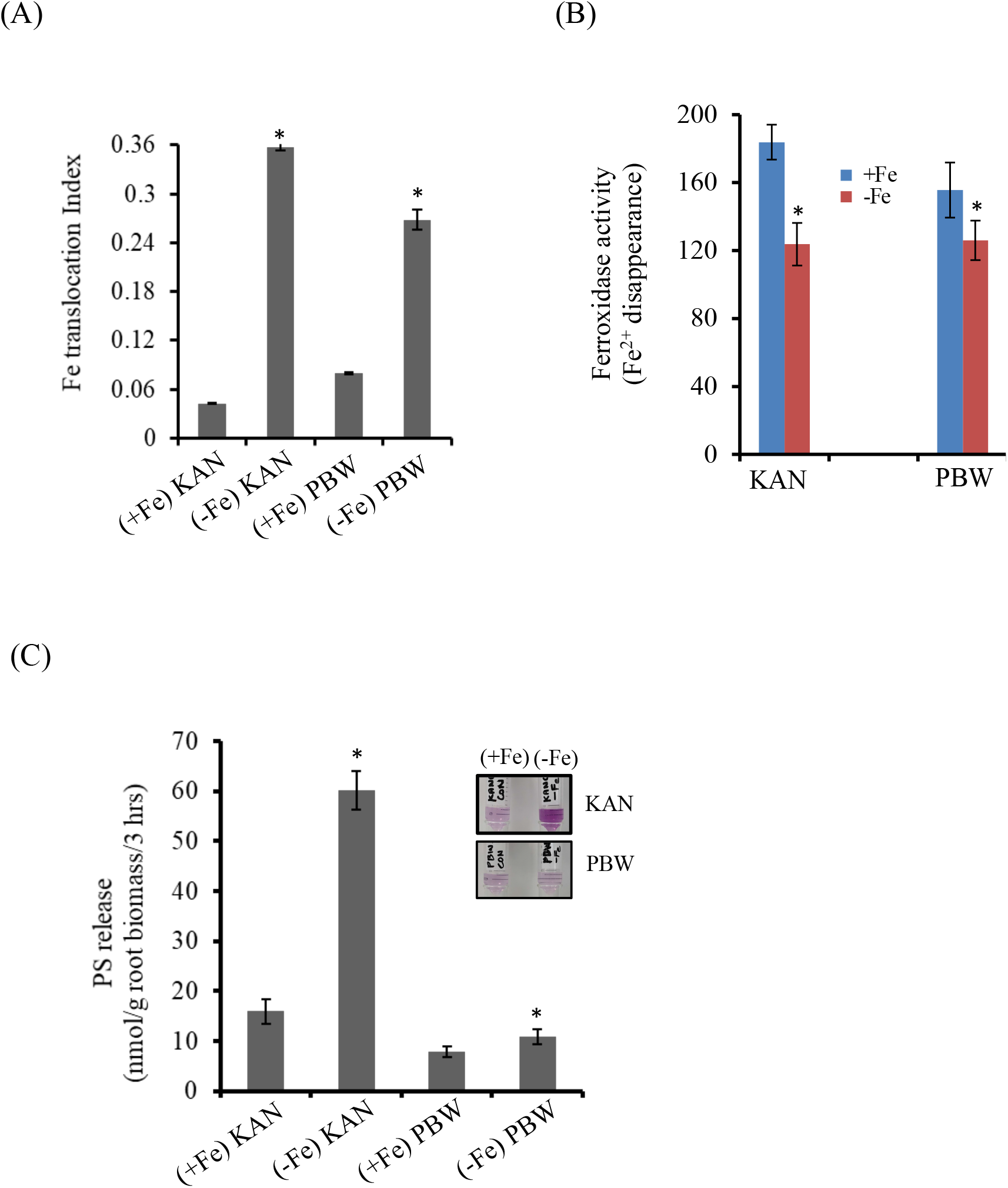
Effect of Fe deficiency on the translocation in KAN and PBW. (A) Fe translocation index in hexaploid wheat subjected to Fe deficiency. (B) Ferroxidase activity measurements in hexaploid wheat subjected to Fe deficiency. (C) PS released by the roots of the wheat cultivars grown under Fe deficiency and control conditions.

Specifically, in shoots, multiple ZIFL shows high fold expression response in KAN compared to the PBW (Figure 7A). This suggests that at the cellular level, MA or its derivatives could be involved in the mobilization of Fe. Next, we studied the expression levels of the efflux of PS identified as TOM in roots. Our expression studies indicated a high expression of the TOM1 (ZIFL4.1/ZIFL4.2) in the KAN compared to the PBW genotype (Figure 7A&B). Overall, these data indicate that chlorotic response is transduced to the roots to initiate the process of Fe deficiency adaptation that help in its remobilization or enhancing its translocation. Based on the data, we speculated that the high PS released by KAN account for the high uptake of Fe in roots mobilized to the shoots by the involvement Fe-NA complex. To check the response of genes involved in NA biosynthesis. Our analysis revealed that multiple NAS encoding genes involved in the biosynthesis of NA were upregulated in KAN, whereas only a few NAS was upregulated in PBW (Supplementary Table S5 and Figure 7C). This suggested an enhanced processing activity of the NAS proteins to generate the NA.

**Figure 7:**
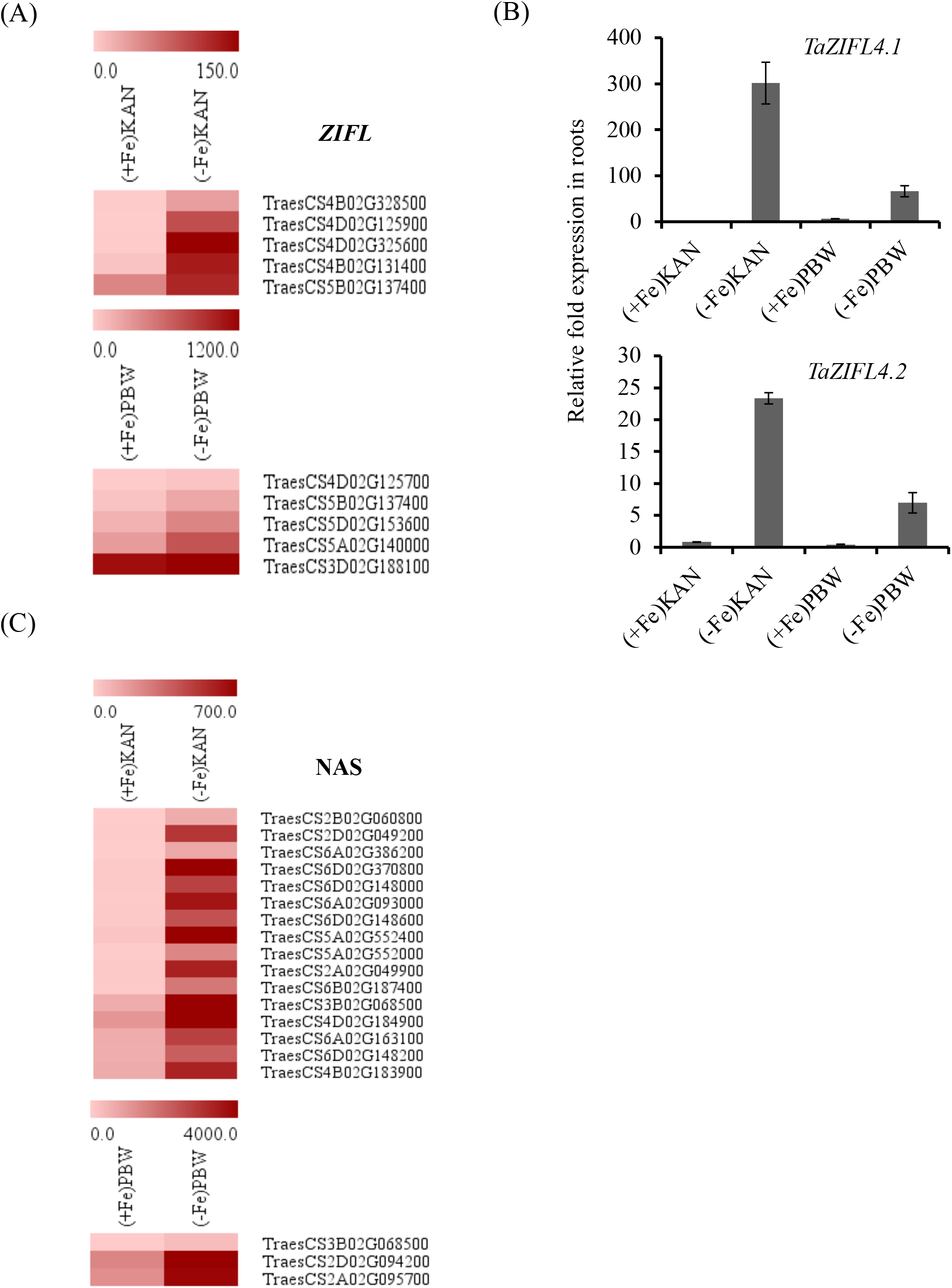
Differential expression of ZIFL and NAS encoding genes in KAN and PBW under Fe deficiency. (A) Heatmap expression of ZIFL transcript in wheat cultivars under Fe deficiency and control condition. (B) qRT-PCR analysis of ZIFL4.1 and ZIFL4.2 genes in the root tissues of KAN and PBW. The C_t_ values were normalized against the wheat ARF1 as internal control. Vertical bars indicate the standard deviation. Sign (*) on the bar indicates that the mean is significantly different at p<0.05 with respect to the control. (C) Heatmap expression of multiple NAS transcript in wheat cultivars under Fe deficiency and control condition.

## Discussion

Previously, we observed the expected conserved gene expression response in different wheat cultivars when exposed to Fe deficient conditions [28,34]. In the current study, molecular evidence was linked to the better tolerance of hexaploid wheat KAN for Fe deficiency. Our study provides multiple datasets that provide a mechanistic insight into how the molecular machinery is recruited to delay the negative impact of fluctuating Fe availability conditions. This work and previous observations suggest that genetic variability is prevalent in the wheat genotype to contribute towards the Fe deficiency at the rhizospheric level. This variability was also linked with the physiological changes in the shoots and chlorophyll biosynthesis in shoots [15].

The transcriptomics-based approach provides the means to address the genetic and molecular basis for the genotypic variation occurring between the cultivars for Fe deficiency. This study led to the confirmation of observations that in susceptible cultivars, the Fe deficiency sensing by the roots is quickly conveyed to the shoots, and this process is delayed in the Fe deficiency tolerant cultivar. Our data suggest that KAN could withstand the Fe deficiency shock and withstand the changes better than the PBW. Specifically, PBW shows the quick and higher intensity transcriptomics for genes involved in photosystem functioning changes compared to KAN (Figure 4). This provides the clue that the susceptible cultivar was highly impacted by the nutrient changes in the roots. In contrast, the KAN shows robust and high differential expression of genes involved in Fe-homeostasis, especially in Fe-uptake and translocation.

Multiple genes have been reported in hexaploid wheat specifically involved in Fe uptake in roots and ion translocation in shoots [28,34]. The ability of KAN to survive Fe deficient conditions was reflected by high PS release, high Fe-mobilization capability, and delayed chlorophyll degradation by enhancing the translocation of Fe to the shoots. The inability of the susceptible cultivar could be linked to the low Fe mobilization to the shoots. Secretion of PS helps mobilise Fe forms, specifically (Fe3+) that is mediated by well-known YSL transporters [35,36]. The secretion of these PS is usually carried out by TOM effluxers [37]. Earlier multiple wheat ZIFL proteins speculated to be the candidate TOM were reported[24,38]. In our study, candidate TOM1 (encoded by ZIFL4.1/4.2) was shown to be highly upregulated (Figure 7A&B) in the KAN cultivar compared to PBW. Interestingly, only ZIFL4.1 and 4.2 were up-regulated in both cultivars at different fold levels. ZIFL4 genes were also shown to be up-regulated during Fe deficiency in wheat cultivar C-306 [28]. This could be interesting since it is possible that the wheat TOM1 could be the functional ortholog of rice TOM1 (OsZIFL4). Earlier, PS release on the root tips was reported to be higher in hexaploid than durum wheat [39]. Genome complexity contributes to the Fe deficiency response trait, as noted earlier by multiple studies [28,34]. It needs to be further investigated if some genomic-biasness interferes with the PS release ability in these genotypes.

Translocation of Fe in the shoots is mainly accorded to the functioning of NA and citrate levels [3]. NA is the major player, and it’s synthesized by using S-adenosyl methionine mediated by NAS proteins [3]. We observed that KAN showed a very high number (13) of transcripts encoding for NAS genes, whereas a low count (3) for the transcript was noted for PBW. Previously, genome-wide characterization of NAS was done and was shown to be differentially expressed under Fe-deficient conditions [40]. Based on that annotation, in our study specifically, *TaNAS2, TaNAS3, TaNAS4, TaNAS7* were highly induced in the tolerant cultivar. High accumulation of the NAS transcripts could account for the enhanced biosynthesis of NA in the shoot which could directly affect the Fe-translocation. In rice, overexpression of barley NAS protein could result in high Fe sequestration and translocation [41,42]. Although speculative, it may hold true that the higher level of NA in KAN shoots could contribute to efficient Fe-mobilization.

Fe deficiency largely alters the functioning of the entire photosynthetic apparatus, including PS1, PSII, and Ferredoxin status and its connection to the Calvin cycle [45,46]. Monocots such as barley was reported to be Fe-deficiency tolerant, employing optimizing the use of PSII downstream complex [47]. The ability to bear the Fe deficiency should be reflected by the minimal fluctuation in the expression of the transcripts involved in photosynthesis. In the susceptible cultivar PBW, we see high DEGs related to PSII system that affects light-harvesting complex-PSII, ferredoxins, state transition and members of the Calvin cycle (Figure 4). Gene expression analysis and the measurement of SPAD values in the leaves suggest that minimal variation was observed for these photosystem pathway genes in the cv. KAN. This reflects that KAN could bear the fluctuation caused by the Fe deficiency at this time compared to PBW.

The current study identifies the molecular determinants of Fe-deficiency tolerance in hexaploid wheat (Figure 8). Our working model reveals multiple characteristics that could account for the Fe deficiency tolerance, including high PS release coupled with the efficient translocation of Fe and high biochemical activity. Our work justifies that KAN could tolerate or delay the early Fe-deficiency symptoms in shoots. This notion was supported by our molecular and biochemical data. Addressing such variability among the hexaploid genotypes of bread wheat will contribute to our ability to generate new germplasm that could tolerate the Fe deficiency condition in soils with minimal productivity and nutritional quality losses.

**Figure 8:**
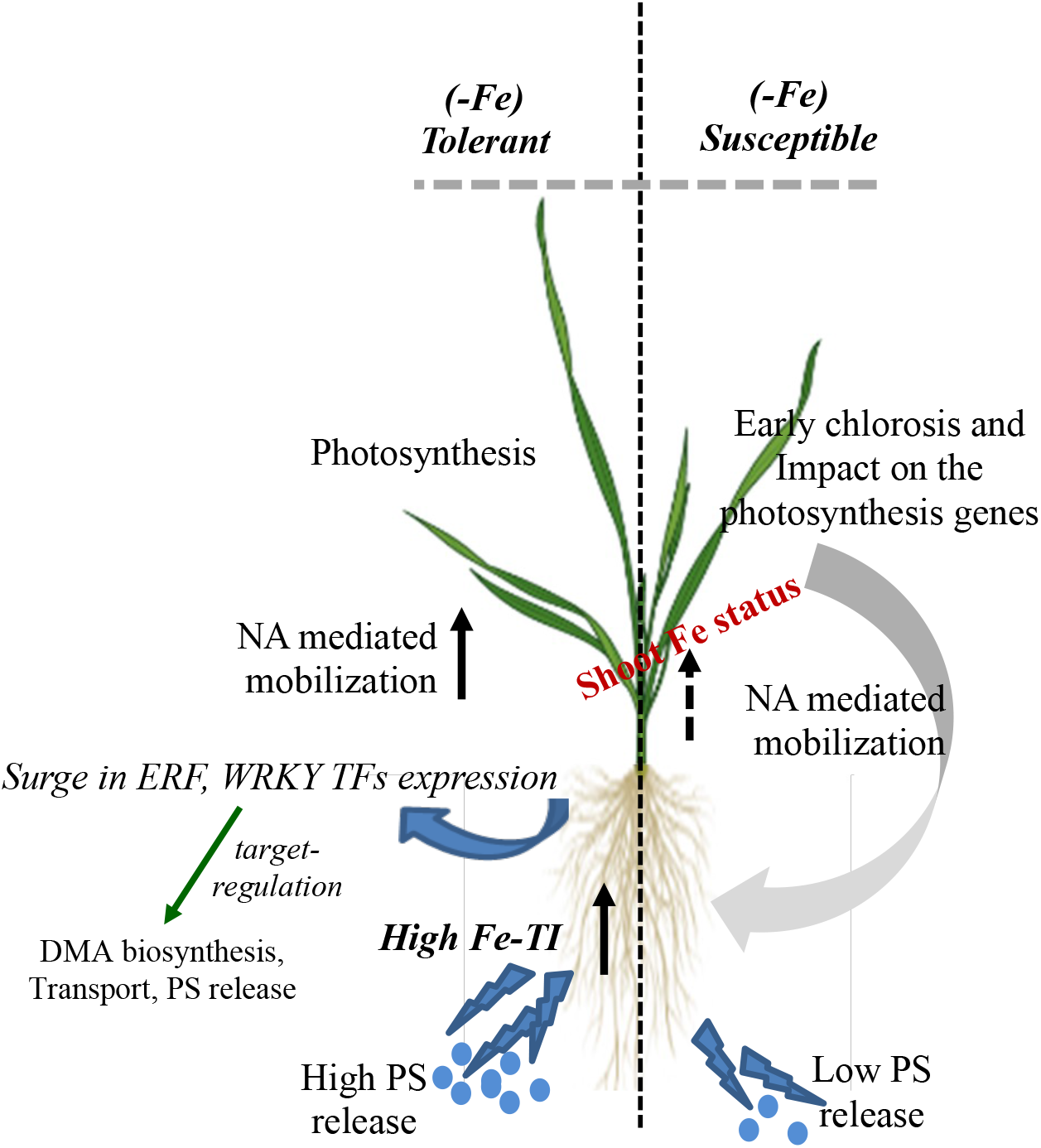
Model describing the molecular and biochemical components involved in imparting tolerance to the Fe deficiency in hexaploid wheat. The contrasting wheat cultivars could differ in the Fe translocation index, enhanced expression of the PS biosynthesis genes and its release in roots, higher expression of NAS genes to mobilize shoot Fe via Nicotianamine and robust photosynthetic response to provide Fe deficiency response. The compounding response of these activities was observed in the tolerant wheat cultivars as compared to Fe sensitive cultivar.

## Abbreviation

KAN: Kanchan
PBW: PBW-343
NAS: Nicotianamine synthase
NA: nicotianamine
Fe: Iron
PS: Phytosiderophores
MDA: Malondialdehyde.

## Acknowledgements

The authors thank Executive Director, NABI for facilities and support. This work was funded by the NABI-CORE grant to AKP. VM acknowledge fellowship support from UGC-CSIR. Part of this work was also supported by the Newton Bhabha PhD Placement Program 2020 granted to VM. DBT-eLibrary Consortium (DeLCON) is acknowledged for providing timely support and access to e-resources for this work. The wheat genome resources developed by International Wheat Genome Sequencing Consortium are highly appreciated.

## Author contributions

Conceptualization: AKP, VM and BS; Data curation: GK, VM, AKP; Formal analysis: VM, BS, GK, JKR, PS and RJ; Funding acquisition: AKP and JKR; Investigation: VM, GK, PC, DT and AKP; Methodology: VM, AS, BS, PC, AM; Project administration: AKP and BS; Resources: JR, AKP, RJ; Supervision: AKP, BS and JKR; Roles/Writing - original draft: AKP, VM and GK; Writing - review & editing: JKR, RJ and DT.

